# Distinct and overlapping functions of *Miscanthus sinensis* MYB transcription factors SCM1 and MYB103 in lignin biosynthesis

**DOI:** 10.1101/629709

**Authors:** Philippe Golfier, Faride Unda, Emily K. Murphy, Jianbo Xie, Feng He, Wan Zhang, Shawn D. Mansfield, Thomas Rausch, Sebastian Wolf

## Abstract

Cell wall recalcitrance is a major constraint for the exploitation of lignocellulosic biomass as renewable resource for energy and bio-based products. Transcriptional regulators of the lignin biosynthetic pathway represent promising targets for tailoring lignin content and composition in plant secondary cell walls. A wealth of research in model organisms has revealed that transcriptional regulation of secondary cell wall formation is orchestrated by a hierarchical transcription factor (TF) network with NAC TFs as master regulators and MYB factors in the lower tier regulators. However, knowledge about the transcriptional regulation of lignin biosynthesis in lignocellulosic feedstocks, such as Miscanthus, is limited. Here, we characterized two Miscanthus MYB TFs, MsSCM1 and MsMYB103, and compared their transcriptional impact with that of the master regulator MsSND1. In Miscanthus leaves *MsSCM1* and *MsMYB103* are expressed at growth stages associated with lignification. Ectopic expression of *MsSCM1* and *MsMYB103* in tobacco leaves was sufficient to trigger secondary cell wall deposition with distinct sugar and lignin composition. Moreover, RNA-seq analysis revealed that the transcriptional responses to *MsSCM1* and *MsMYB103* overexpression showed extensive overlap with the response to *MsSND1*, but were distinct from each other, underscoring the inherent complexity of secondary cell wall formation. Together, *MsSCM1* and *MsMYB103* represent interesting targets for manipulations of lignin content and composition in Miscanthus towards tailored biomass.

## Introduction

Lignocellulosic biomass, which largely exists in the form of plant secondary cell walls (SCW), holds enormous potential as renewable feedstock for a sustainable economy (Pauly and Keegstra, 2010). Despite various biorefining applications, processing of lignocellulosic biomass into bio-based products and energy is still hampered by the inherent resistance of cell walls to deconstruction, which is largely conferred by lignin (Himmel *et al*., 2007; Pauly and Keegstra, 2010). The aromatic polyphenol lignin associates with the cellulose and hemicellulose network of SCWs, providing mechanical support, rigidity, and hydrophobicity. The mechanical importance of lignin is highlighted by lignin-deficient mutants that manifest in dwarfism, collapsed xylem vessels, or higher susceptibility against pathogens, although some of those phenotypes can also be the result of signalling-mediated secondary responses (Bonawitz and Chapple, 2013). Due to cell wall recalcitrance, lignin engineering has become a focal point of efforts aiming to alter lignin content, composition, and structure, exploiting the inherent natural plasticity of lignin (Mottiar *et al*., 2016). Thus, it is vital to understand lignin biosynthesis and its regulation to harness the potential of lignocellulosic biomass as a renewable resource for biorefineries.

The genus Miscanthus is regarded as one of the most promising energy crops for the production of lignocellulosic biomass in temperate climates. Miscanthus species are rhizomatous, perennial grasses that combine efficient C4 metabolism and favorable leaf morphology resulting in high yields with modest water and nutrient requirement. Breeding programs have generated novel Miscanthus hybrids to overcome major limitations for adoption into diverse agricultural systems displaying different soil qualities and wide climatic ranges (Clifton-Brown *et al*., 2016). Moreover, Miscanthus breeding has focused on the development of molecular markers to facilitate selection of varieties with improved biomass yield and/or composition (da Costa *et al*., 2017). Genetic resources for Miscanthus, such as an extensive transcriptome database (Barling *et al*., 2013) and a recently completed *Miscanthus sinensis* draft genome (*Miscanthus sinensis* v7.1 DOE-JGI, http://phytozome.jgi.doe.gov/), are now available to explore biological processes with molecular approaches. A detailed, mechanistic understanding of SCW formation has the potential to complement marker-assisted breeding of novel Miscanthus traits with tailored biomass.

The irreversibility of SCW formation requires a tight regulation of the biosynthesis of cellulose, hemicellulose, and lignin. A sophisticated multi-tiered network of NAM, ATAF1,2, CUC (NAC), and MYB transcription factors (TFs) coordinately integrate developmental and environmental aspects (Nakano *et al*., 2015; Zhong and Ye, 2015; Taylor-Teeples *et al*., 2015; Houston *et al*., 2016). In Arabidopsis, the closely related NAC SECONDARY CELL WALL THICKENING PROMOTING FACTOR (NST)1 and 2, SECONDARY WALL-ASSOCIATED NAC DOAMAIN1 (SND1)/NST3, and VASCULAR-RELATED NAC DOMAIN (VND) 6 and 7 have been identified as master switches regulating SCW formation, including regulation of biosynthetic pathways for cellulose, hemicellulose, and lignin (Kubo *et al*., 2005; Zhong *et al*., 2006, 2007*a*; Mitsuda *et al*., 2007; Nakano *et al*., 2015), as well as controlling the patterned deposition of SCW components (Kubo *et al*., 2005). Downstream of NAC TFs, MYB46 and MYB83 are situated as second tier regulators, and were shown to act redundantly during SCW formation (Zhong *et al*., 2007*b*; McCarthy *et al*., 2009). A large number of lower tier members such as MYB20, MYB42, MYB43, MYB58, MYB63, MYB83, MYB85, and MYB103, are often targets of both the second tier MYB46 and 83 as well as of NAC master switches, and are assumed to transcriptionally fine-tune SCW biosynthesis, particularly lignin biosynthesis (Zhong *et al*., 2008; Zhou *et al*., 2009; Zhao and Dixon, 2011). However, lower tier MYB factors from grasses seem to regulate additional secondary cell wall components along with lignin (Noda *et al*., 2015; Scully *et al*., 2016). For example, MYB103 in Arabidopsis has been described to specifically affect lignin composition without affecting total lignin and cellulose content (Öhman *et al*., 2013). However, MYB103 orthologues from grass species seem to regulate a much broader spectrum of SCW-related target genes (Hirano *et al*., 2013; Yang *et al*., 2014; Ye *et al*., 2015). Thus, while the SCW transcriptional network seems to be conserved to some degree, recent evidence suggests grass-specific nuances exist (Rao and Dixon, 2018), in line with an expansion of the MYB class of TFs in monocots (Rabinowicz *et al*., 1999; Zhao and Bartley, 2014). Recently, MsSND1 and MsSCM1 from *Miscanthus sinensis* have been identified as positive regulators of lignin biosynthesis (Golfier *et al*., 2017). However, a detailed analysis of the regulatory properties of MsSND1 and MsSCM1, as well as the impact of these TFs on SCW composition has not been addressed yet. In addition, it is unclear whether Miscanthus lower tier MYB factors are largely redundant or orchestrate a distinct transcriptional response, which would make them interesting targets for tailoring lignin content and/or composition.

Here, we compare two MYB TFs, namely MsSCM1 (Golfier *et al*., 2017) and the newly identified MsMYB103 with MsSND1, a master switch of SCW formation. In Miscanthus, *MsSCM1* and *MsMYB103* were expressed in tissue undergoing lignification. Ectopically expressed in *N. benthamiana* leaves, *MsSND1, MsSCM1*, and *MsMYB103* were capable of inducing lignin deposition leading to distinct cell wall compositions. A global expression analysis uncovered the transcriptional responses to *MsSND1, MsSCM1*, and *MsMYB103* expression, demonstrating overlapping and diverging underlying cell wall profiles. Taken together, our results suggest MsSCM1 and MsMYB103 act as regulators of lignin biosynthesis leading to distinct lignin qualities.

## Materials and methods

### Plasmid Construction

For this study, plasmid constructs were generated via GreenGate cloning (Lampropoulos *et al*., 2013). The protein-coding region of *MsSND1, MsSCM1*, and *MsMYB103* were amplified by PCR using appropriate primers with BsaI restrictions site overhang and cDNA from *Miscanthus sinensis* (Identification code Sin-13; Clifton-Brown and Lewandowski, 2002). More information about primer sequences, GreenGate modules, and assembled constructs for ectopic expression can be found in Supplementary Tab. 1.

### Gene expression analysis

Gene expression analysis was performed according to Golfier *et al*., (2017). Briefly, total RNA was extracted from 30 mg ground leaf tissue using GeneMatrix Universal RNA Purification Kit (EURx/Roboklon). 1 μg of RNA was reverse transcribed using oligo dT primer either by Superscript III Reverse Transcriptase (Invitrogen) for tobacco or by RevertAid First Strand cDNA Synthesis Kit (Thermo Fisher) for Miscanthus according to manufacturer’s instructions. The quantitative Real-Time PCR was carried out in a total volume of 15 μl containing 2 μl of diluted cDNA, 1 μl 5 μM forward and reverse primer (Supplementary Tab. 1.), 0.3 μl of each 10 mM dNTPs, 1:400 diluted SYBR(R) Green I (Sigma-Aldrich), JumpStart(TM) Taq DNA polymerase in the corresponding buffer, and rnase-free H_2_O. Measurements were observed with a Rotor-Gene Q thermocycler and evaluated by Q Series Software Q (Qiagen). Gene expression was normalized against UBC and PP2A in Miscanthus and PP2A in tobacco.

### Nicotiana Benthamiana Infiltration

For ectopic expression of TF, the respective GreenGate construct was transformed into *Agrobacterium tumefaciens* ASE (pSOUP^+^). Selected transgenic clones were incubated in liquid LB medium containing respective antibiotics for two days. The LB medium was replaced with infiltration medium (10 mM MgCl_2_, 10 mM MES, 0.15 mM acetosyringone at pH 5.7). Leaves of 4-6 weeks-old *Nicotiana benthamiana* were co-infiltrated with bacteria suspension of transgenic clones and *A. tumefaciens* C58C1, containing a p14 silencing suppressor (Merai *et al*., 2005). An empty vector control (Supplementary Tab. 1) was co-infiltrated with p14 Agrobacterium as control.

### Tissue Staining and Microscopy

After 5 days of ectopic expression, *N. benthamiana* leaves were embedded in 6% agarose and hand-sectioned with a razor blade. For Basic Fuchsin staining, cross-sections were cleared and fixed for 15 min in methanol, before they were incubated in 10% (w/v) NaOH at 65 °C for 1 hour. Lignin was stained with 0.01% (w/v) Basic Fuchsin in water for five minutes and washed briefly with 70% (v/v) ethanol. Cross-sections were mounted in 50% (v/v) glycerol and imaged on a confocal Leica TCS SP5II microscope equipped with a 40.0 × 1.25 NA objective under 561 nm excitation and 593/40 nm emission. ImageJ software was used to produce orthogonal sections and scale bars.

### Cell Wall Analysis

*N. benthamiana* leaves were harvested 5 days post infiltration. Non-infiltrated plant tissue like midribs and mature veins were removed and dried for 2 days at 40°C. Dry plant material was ground with a Wiley mill and subsequently Soxhlet extracted with hot acetone (70°C) for 24 h. Extractive-free plant material was used to determine Klason lignin content and structural carbohydrate concentration according to Coleman *et al*., 2008. The monolignol composition was determined by thioacidolysis as described in Robinson and Mansfield (2009) with minor modifications. Since the overall lignin content was relatively low, lignin monomers were extracted from 20 mg of extractive-free plant tissue and 2 μl were injected into the gas chromatograph.

### Transcriptome Analysis

To minimize the effects of natural transcriptional differences between *N. benthamiana* plants, single leaves were infiltrated with all four TFs according to the scheme illustrated in Supplementary Fig. S2. In order to capture high gene expression of transcriptional networks, leaf tissue was harvested 4 days post infiltration and immediately frozen and ground in liquid nitrogen. Total RNA was extracted from 50 mg ground leaf tissue using GeneMatrix Universal RNA Purification Kit (EURx/Roboklon). Library preparation was performed by GATC Biotech (Konstanz, Germany) using proprietary methods. Subsequently, the strand-specific cDNA library was sequenced by GATC Biotech on Illumina HiSeq 4000 instruments in 150 bp paired-end mode.

The re-annotated *N. benthamiana* transcriptome (Kourelis *et al*., 2018) was downloaded from ORA (https://ora.ox.ac.uk/objects/uuid:f09e1d98-f0f1-4560-aed4-a5147bc7739d). The index of the reference transcriptome was built using Bowtie2 (Langmead and Salzberg, 2012) and paired-end RNA sequencing reads were aligned to the reference transcriptome using RSEM (Li and Dewey, 2011) on Galaxy (RSEM-Bowtie2 v0.9.0). Differentially expressed genes (DEGs) between control- and TF-infiltration lines were identified with edgeR using read counts for each transcript and TMM normalization. Genes with an adjusted p-value <0.05 and a minimum two-fold change were considered as differentially expressed (Supplementary Tab. 2.). The heatmap was visualized with normalized counts (logCPM) that were hierarchically clustered by average linkage and one minus Pearson correlation of rows using Morpheus (https://software.broadinstitute.org/morpheus). GO enrichment was performed based on the hypergeometric test, and P value was corrected using FDR method.

### Accession numbers

AmMYB330 (P81393), AtMYB20 (AT1G66230.1), AtMYB43 (AT5G16600.1), AtMYB46 (AT5G12870.1), AtMYB58 (AT1G16490.1), AtMYB61 (AT1G09540.1), AtMYB63 (AT1G79180.1), AtMYB83 (AT3G08500.1), AtMYB85 (AT4G22680.1), AtMYB103 (AT1G63910.1), BdSWAM1 (KQK09372), EgMYB1 (CAE09058), EgMYB2 (CAE09057), GhMYBL1 (KF430216), MsSCM1 (KY930621), MsSCM2 (MF996502), MsSCM3 (KY930622), MsSCM4 (MF996501), MsMYB103 (MK704407), OsMYB103L (LOC_Os08g05520.1), PbrMYB169 (MG594365), PtMYB1 (AAQ62541), PtMYB4 (AAQ62540), PtMYB8 (ABD60280), PtoMYB92 (KP710214), PtrMYB2 (AGT02397), PtrMYB3 (AGT02395), PtrMYB10 (XP_002298014), PtrMYB20 (AGT02396), PtrMYB21 (AGT02398), PtrMYB128 (XP_002304517), PtrMYB152 (POPTR_0017s02850), PvMYB63A (Pavir.Bb02654), PvMYB85A (Pavir.Gb00587), SbMYB60 (Sobic.004G273800).

## Results and Discussion

### Miscanthus MYB103 and SCM1 are phylogentically related to MYB TFs regulating lignin formation

In order to find potential targets in Miscanthus that allows engineering of biomass optimized for bioenergy and biomaterials applications, we searched suitable TFs with the ability to regulate SCW formation with a focus on lignin biosynthesis. In Miscanthus, the NAC TF MsSND1 has been previously characterized as a master regulator orchestrating SCW formation, and MsSCM1, a MYB TF related to MYB20/43/85 (Zhong *et al*., 2008), has been proposed as specific regulator of lignin biosynthesis (Golfier *et al*., 2017). We identified a putative MsMYB103 in a Miscanthus transcriptome (Barling *et al*., 2013) by a homology-based approach with AtMYB103 as query sequence. AtMYB103 has been suggested as a key gene regulating the expression of ferulate-5-hydroxylase (F5H), which directs lignin biosynthesis towards formation of syringyl-rich lignin (Öhman *et al*., 2013) and was initially suggested to regulate lignin composition exclusively. However, MYB103 orthologues of grass species have also been shown to promote cellulose and hemicellulose genes (Hirano *et al*., 2013; Yang *et al*., 2014; Ye *et al*., 2015) and AtMYB103 at least associates with cellulose synthase promoter sequences (Zhong *et al*., 2008; Taylor-Teeples *et al*., 2015). The predicted MsMYB103 protein shares 43.9% identity and 51.9% similarity on amino acid level with AtMYB103 (Supplementary Fig. S). An alignment of the R2R3 MYB DNA-binding domain at the N-terminus showed 96% identity on amino acid level, suggesting similar DNA binding characteristics between MsMYB103 and AtMYB103. Phylogenetic analyses revealed a close relationship of MsMYB103 to AtMYB103, GhMYBL1, PtrMYB10, and PtrMYB128 (Fig. 1), TFs known to participate in the regulation of lignin biosynthesis and SCW formation (Zhong *et al*., 2011; Öhman *et al*., 2013; Sun *et al*., 2015). Similarly, MsSCM1 falls into a clade of SCW regulators including AtMYB20/43/85, PbrMYB169, PtoMYB92, and PvMYB85 (Fig. 1; Zhong *et al*., 2008; Li *et al*., 2015; Rao *et al*., 2018; Xue *et al*., 2019).

**Fig. 1.**
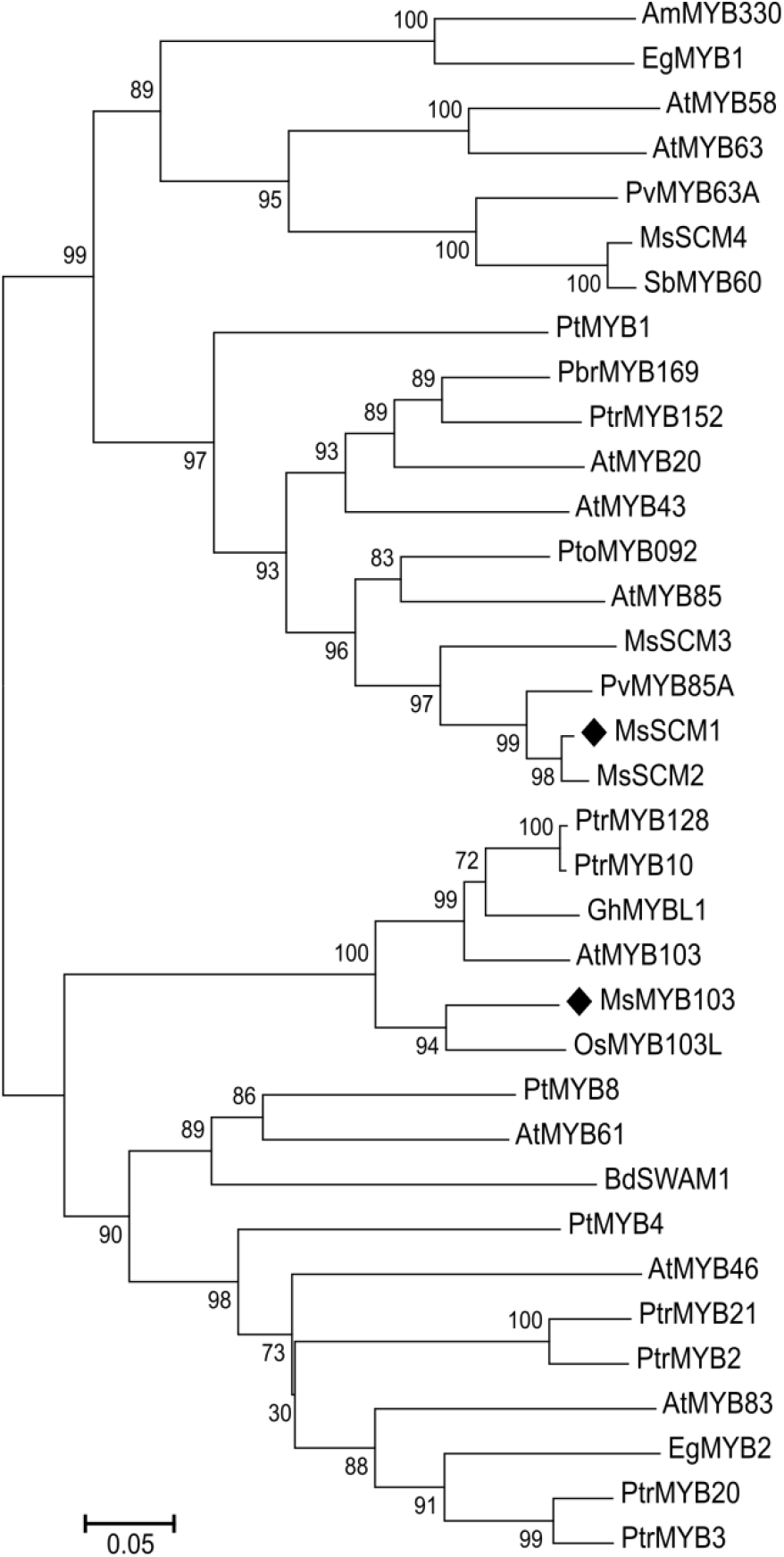
Phylogenetic analysis of MsSCM1-4 and MsMYB103 in relation to MYB transcription factors involved in regulating phenylpropanoid and lignin biosynthesis from angiosperm lineages *Arabidopsis, Antirrhinum, Brachypodium, Eucalyptus, Gossypium, Oryza, Panicum, Pinus, Populus, Pyrus*, and *Sorghum*. Amino acid sequences were aligned by ClustalW and the phylogenetic tree was calculated by MEGA6 software with neighbor-joining with 1000 bootstraps (Tamura *et al*., 2013). The evolutionary distance is represented by the scale bar.

### Expression of *MsSCM1* and *MsMYB103* is associated with tissues undergoing lignification

In monocots, leaf growth is controlled by an intercalary meristem that is situated at the stem node, driving leaf elongation. The continued differentiation along the leaf axis from immature leaf sheath towards mature leaf tip results in a linear developmental gradient. Previously, high expression of *MsSND1* was found to coincide with vascular differentiation and SCW formation in Miscanthus leaves (Golfier *et al*., 2017), implying a role for MsSND1 in those processes. To explore the potential involvement of *MsMYB103* and *MsSCM1* in secondary cell wall formation in Miscanthus, expression of *MsMYB103* and *MsSCM1* was determined together with *MsSND1* along the leaf gradient and visualized in a heat map (Fig. 2). The transcript abundance of *MsSND1* was highest at the leaf base in the first segment, and decreased strongly in the following segments, confirming previous findings (Golfier *et al*., 2017). The expression of *MsMYB103* and *MsSCM1* reached its maximum in the second segment and declined markedly in the following segments (Fig. 2). Putative orthologues of *MsMYB103* and *MsSCM1* in rice and maize (GRMZM2G325907 and LOC_Os08g05520, as well as GRMZM2G104551 and LOC_Os09g36250, respectively) display a comparable expression pattern (Wang *et al*., 2014) in immature leaf, an organ known to show active SCW formation and lignification. This is reminiscent of other putative *MsSND1* target genes (Li *et al*., 2010; Golfier *et al*., 2017) and supports a role of *MsMYB103* and *MsSCM1* in SCW formation and lignification.

**Fig. 2.**
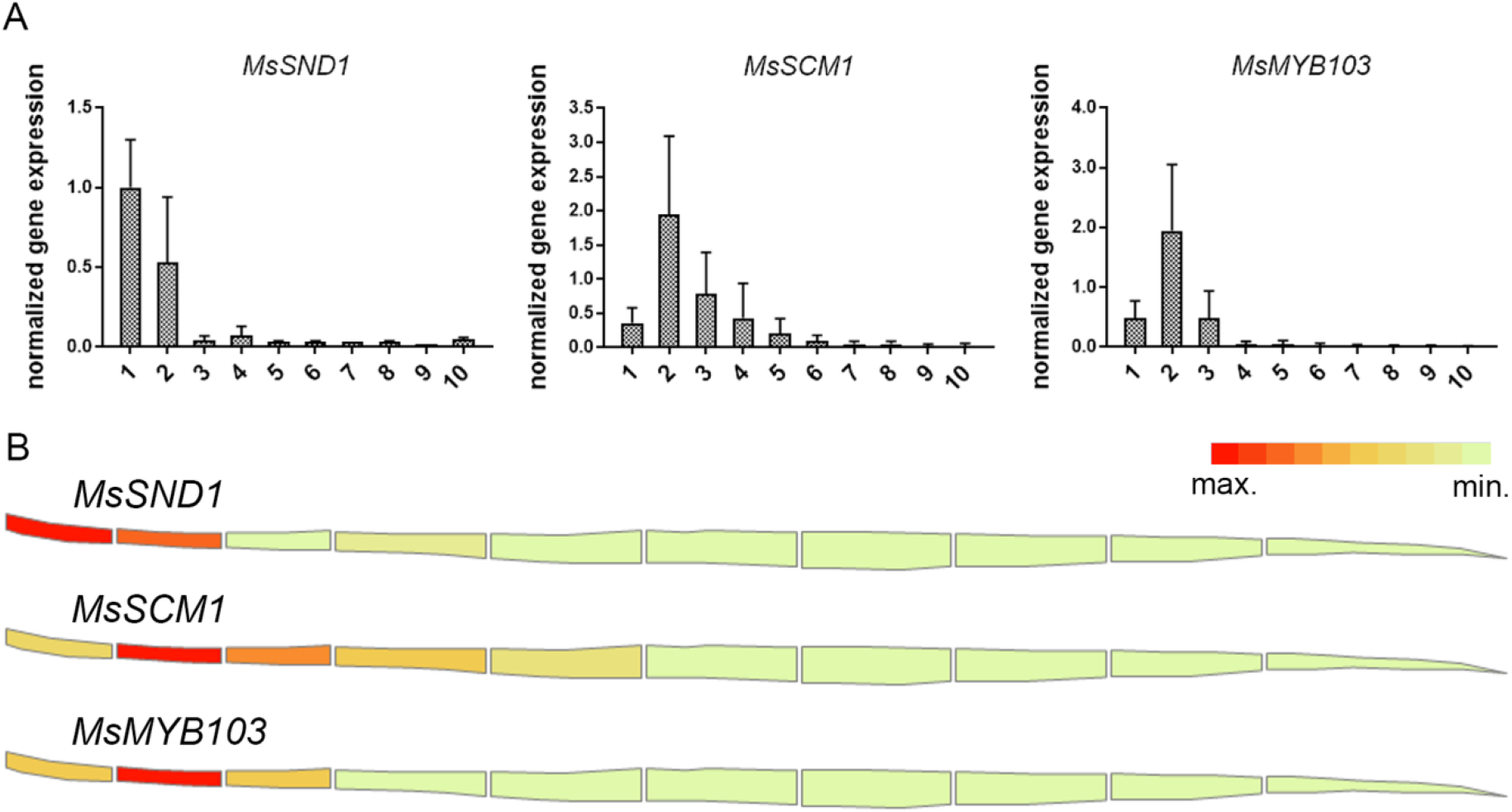
Expression of *MsSND1, MsSCM1*, and *MsMYB103* is associated with tissues undergoing lignification. Leaves were dissected into ten sections spanning from immature leaf base at the right side to mature leaf tip on the left side. Relative gene expression values are calculated from three biological replicates which are normalized against two reference genes (UBC and PP2A) and visualized as a heat map.

### Ectopic expression of *MsSCM1* and *MsMYB103* lead to lignin deposition in tobacco leaves

In order to investigate the capability of *MsSND1, MsSCM1*, and *MsMYB103* to induce lignification, these TFs were ectopically expressed in *Nicotiana benthamiana* leaves. As previously described (Golfier *et al*., 2017), *MsSND1* induced patterned formation of SCWs in spongy mesophyll, epidermal, and palisade cells reminiscent of xylem elements, which were visualized by Basic Fuchsin staining (Fig. 3A). In contrast, ectopic expression of *MsMYB103* resulted in uniform deposition of lignin in most leaf cells and markedly more intense Basic Fuchsin fluorescence in some cells (Fig. 3B), similar to *MsSCM1* (Fig 3C, Golfier *et al*., 2017). This pronounced difference in lignin deposition between *MsSND1* and *MsMYB103/ MsSCM1* expressing cells confirms distinct transcriptional responses to top tier NAC TF and lower tier MYB TF factor expression. It remains elusive why some individual cells show stronger lignification after ectopic expression of MsMYB103/SCM1 than some of their neighboring cells. It is conceivable that only cells with the appropriate cell wall can lignify because incorporation of lignin depends on a suitable polysaccharide matrix (Taylor *et al*., 1992) and appropriate amounts of lignin precursor precursors such as monolignols (Smith *et al*., 2013). In summary, ectopic expression in tobacco leaves revealed that *MsMYB103* and *MsSCM1* are capable of activating the lignin machinery leading to lignin deposition.

**Fig. 3.**
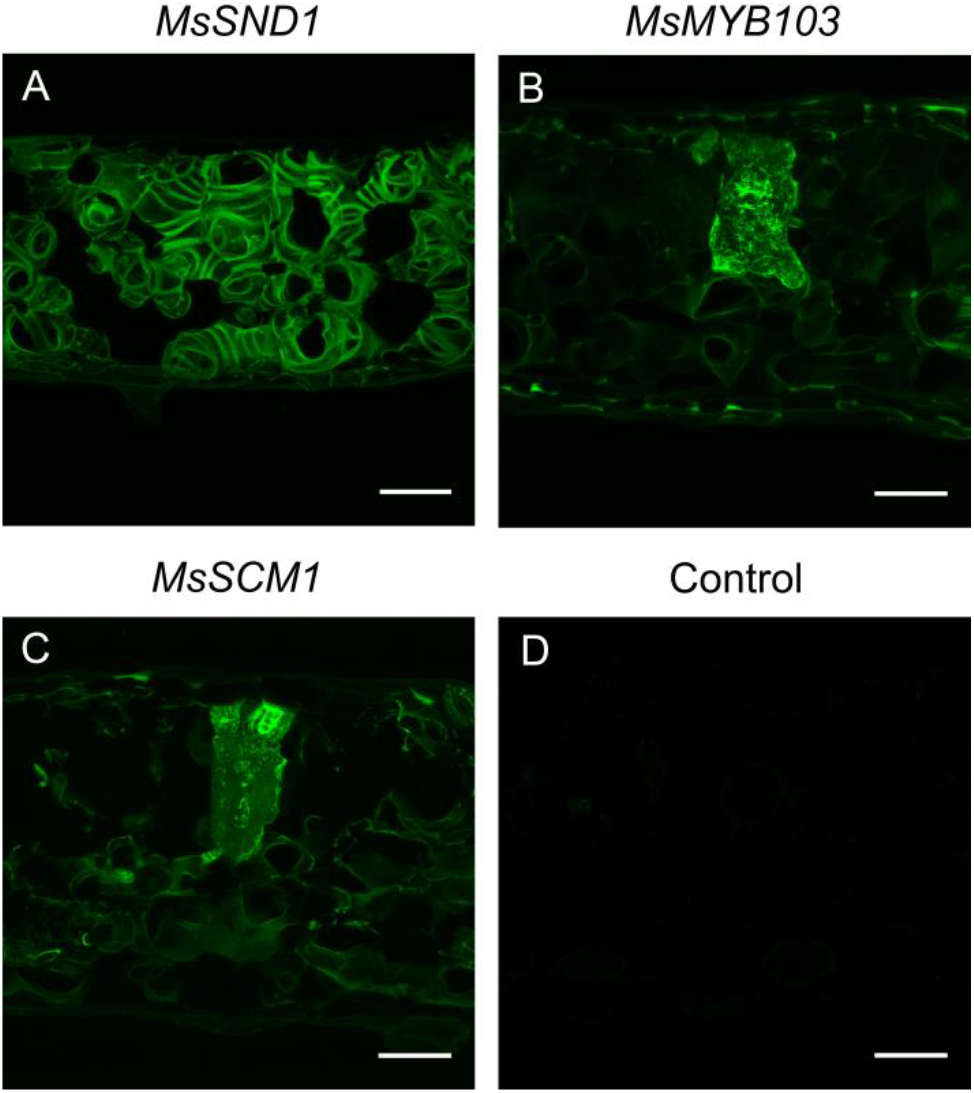
Lignin formation induced by ectopic expression of *MsSND1, MsSCM1*, and *MsMYB103* in *N. benthamiana* leaves. After 5 days of ectopic expression, leaf cross-sections were stained with Basic Fuchsin to observe fluorescence under a confocal microscope. An empty vector control served as control. Scale bar 50 μm.

### Expression of *MsSND1, MsMYB103*, and *MsSCM1* results in distinct secondary cell wall composition

The ability of *MsSND1, MsMYB103*, and *MsSCM1* to regulate cell wall formation prompted us to exploit the tobacco system to investigate compositional changes in the cell walls associated with ectopic expression of the TFs. As expected, in mock-infiltrated tobacco leaves, the content of acid-insoluble Klason lignin was very low (ca. 15 μg/mg dry weight; Fig. 4A). Ectopic expression of *MsSND1, AtSND1, MsSCM1*, and *MsMYB103* elevated Klason lignin content by 2 to 5.5-fold (29 to 82 μg/mg dry tissue). MsSND1 and AtSND1 had a greater effect compared to MsSCM1 and MsMYB103, consistent with the microscopy results obtained with histochemical staining (Fig. 3). Moreover, measurement of non-condensed lignin by thioacidolysis demonstrated a higher S/G ratio in *MsMYB103* expressing tissue, whereas expression of the other transcription factors resulted in lower ratios, similar to the control (Fig. 4B). These results are consistent with the described role of Arabidopsis MYB103 as a key regulator of syringyl lignin (Öhman *et al*., 2013). However, in switchgrass expressing *PvMYB85A*, a close relative of *MsSCM1* (Fig. 1), was capable of elevating the S/G ratio (Rao *et al*., 2019), whereas *MsSCM1* did not change S/G ratio in *N. benthamiana* (Fig. 4B), highlighting functional nuances between two closely related genes. Analysis of the structural carbohydrates revealed that ectopic expression of *MsSND1* and *AtSND1* provoke a strong increase in xylose (Fig. 4C), which likely originates from the formation of xylan and/or arabinoxylan in cell walls (Golfier *et al*., 2017). The composition of structural carbohydrates in MsSCM1 samples was similar to control, except for a reduction in glucose, which was observed in all samples. Interestingly, transient expression of Ms*MYB103* led to a small increase in mannose and xylose (Fig. 4C), possibly suggesting a regulatory role for *MsMYB103* in hemicellulose formation. Taken together, chemical analysis of cell walls after ectopic expression of *MsSND1, MsSCM1*, and *MsMYB103* revealed distinct cell wall compositions.

**Fig. 4.**
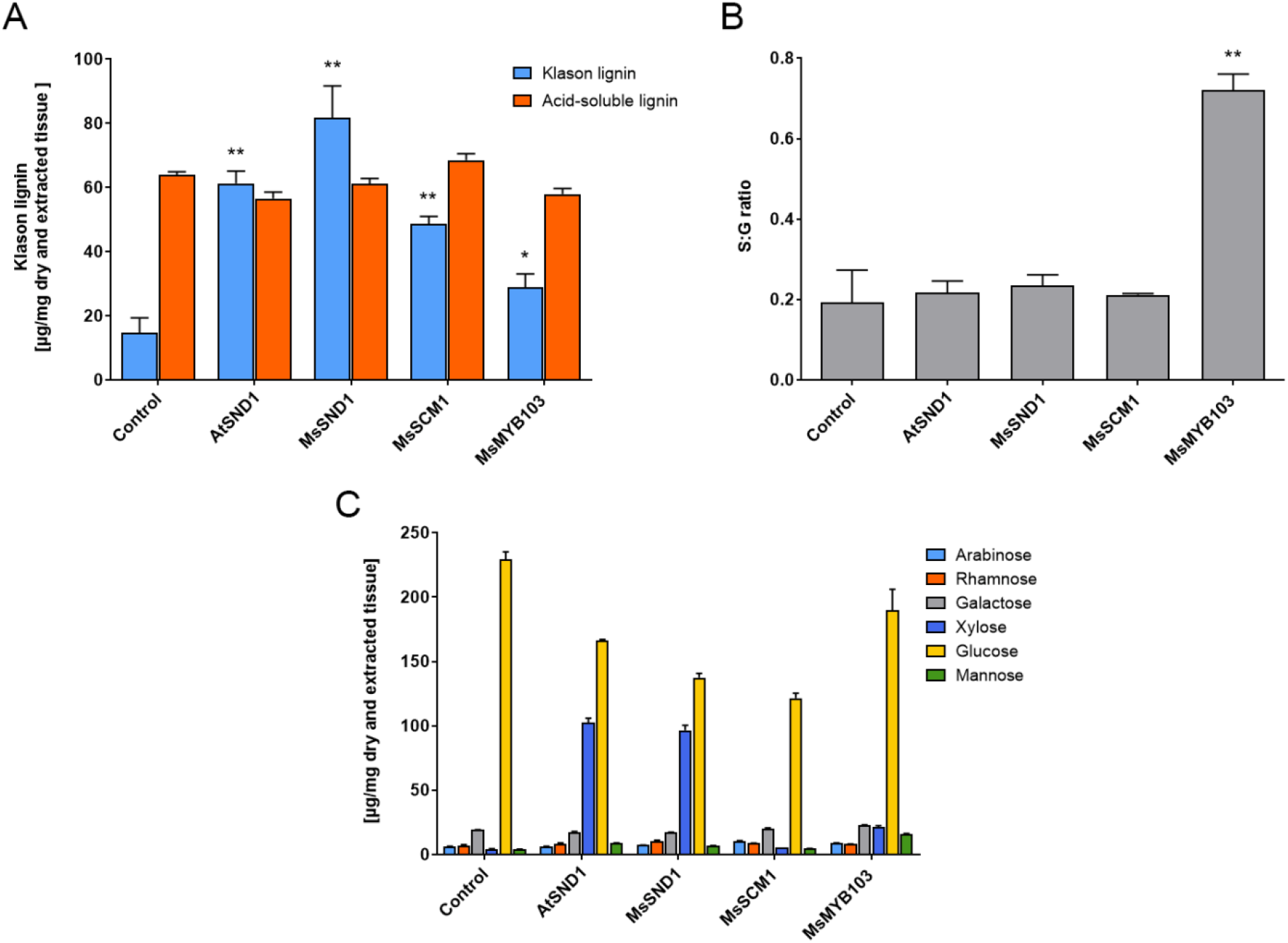
Change in cell wall composition induced by ectopic expression of *MsSND1, MsSCM1*, and *MsMYB103* in *N. benthamiana* leaves. (A) Lignin content was determined by Klason analysis (acid-insoluble lignin) and by OD_205_ (acid-soluble lignin), (B) Ratio of syringyl and guaiacyl monomer-release from thioacidolysis products detected by gas-chromatography, (C) Structural carbohydrate compositions of insoluble extracts were determined by high-pressure liquid chromatography (HPLC), as μg/mg of dry cell walls. As control, an empty vector construct was infiltrated in tobacco leaves. The bars represent mean values from two (B) or three (A, C) biological replicates + SE including two technical replicates for (A, C).

### RNA-seq analysis following expression of *MsSND1, MsSCM1*, and *MsMYB103* delineates diverging transcriptional landscapes

In order to understand how expression of *MsSND1, MsSCM1*, and *MsMYB103* translates into compositional cell wall changes, we pursued profiling of TF-mediated gene expression changes. Agrobacterium-mediated infiltration of TFs into *N. benthamiana* leaves followed by RNA-seq was shown to be suitable to discover downstream targets of TFs (Bond *et al*., 2016). Four days post infiltration, we could verify expression of *MsSND1, MsSCM1, MsMYB103*, and the putative downstream targets *CCoAOMT* and *XCP1* by q-RT-PCR, suggesting that the experimental approach is suitable to capture the transcriptional profiles via RNA-seq (Supplementary Fig. S2). Evaluation of the transcriptomes using stringent parameter revealed a total of 4716 differentially expressed genes (DEGs) in at least one condition compared to the control (Supplementary Tab. 2.). The hierarchically clustered heatmap of normalized counts of DEGs from control, MsSND1, MsSCM1, and MsMYB103 illustrated a very similar expression pattern within biological replicates (Fig. 5A). As expected, the transcriptional response to expression of the master regulator MsSND1 was more extensive (3317 DEGs) than that to MsSCM1 and MsMYB103 expression (2241 and 1392 DEGs, respectively). The samples expressing lower tier MYB factors shared between 47% and 69% of DEGs with MsSND1 samples (Fig. 5B, D), consistent with the described role of MYBs as downstream targets of secondary cell wall NAC master regulators (Rao and Dixon, 2018). However, samples from tissue expressing MsSCM1 shared only 25% of the upregulated DEGs with tissues expressing MsMYB103, whereas the overlap was even lower (16%) with downregulated DEGs (Fig. 5D). This suggests that lower tier MYB factors can have unique targets, possibly contributing diversification and fine-tuning of the secondary cell wall transcriptional program.

**Fig. 5.**
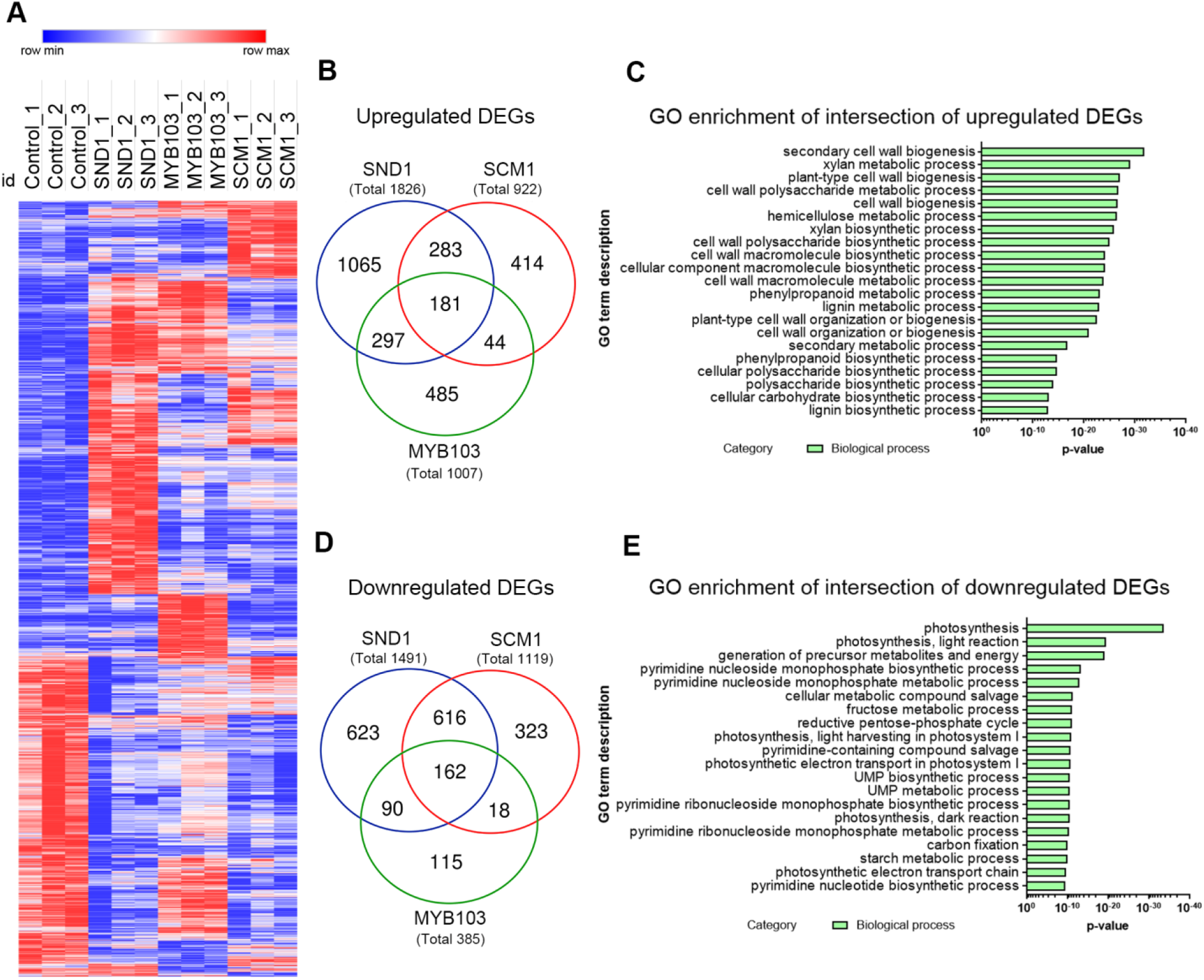
Transcript profiling of transcriptional network regulated by MsSND1, MsSCM1, and MsMYB103. (A) Heatmap of normalized counts (logCPM) of the 4716 genes that were differentially expressed in at least one sample compared to control infiltrations with an empty vector are visualized with Morpheus. Rows were hierarchically clustered by average linkage and one minus Pearson correlation. A detailed heatmap with gene identifier can be found in Supplementary Fig. S3 (B, D) Venn diagram of up- (B) and downregulated (D) DEGs with adjusted p-value < 0.05 and a minimum 2-fold change. (C, E) GO enrichment analysis of the intersection of up- (C) and downregulated (E) DEGs of MsSND1, MsSCM1, and MsMYB103. The first 20 GO term descriptions of the category ‘biological process’ are plotted against the p-value.

Gene ontology (GO) term enrichment analyses of DEGs commonly upregulated by MsSND1, MsSCM1, and MsMYB103 revealed enriched GO terms associated with SCW formation including lignin, xylan, and cellulose biosynthesis (Fig. 5B, C; Supplementary Tab. 3.). In addition, GO terms associated with pectin, in particular homogalacturonan and rhamnogalacturonan I biosynthesis/metabolism, are overrepresented in the transcripts induced by all three TFs (Supplementary Table 3). SCWs are formed following cessation of growth and development often accompany by cell death. GO terms linked to photosynthesis, energy metabolism, and nucleotide metabolism were enriched in the intersection of downregulated DEGs (Fig. 5E), indicating a tightly controlled switch from metabolic active cells towards specialized cells with a role in mechanical support enacted by both top and lower tier TFs in the network. GO enrichment analysis of DEGs suggests distinct and overlapping functions of *MsSND1, MsSCM1*, and *MsMYB103* in lignin biosynthesis and cell wall formation.

### Expression atlas highlights essential genes of lignin biosynthesis

*MsSND1, MsSCM1*, and *MsMYB103* were found to be associated with distinct cell wall and lignin composition (Fig. 4B). To obtain a detailed perspective on genes involved in lignification that may affect lignin quality, expression of putative lignin genes from the transcriptomes were plotted in a schematic lignin biosynthesis pathway (Fig. 6). Ectopic expression of *MsSND1* and *MsSCM1* elevated expression of several putative genes of the general phenylpropanoid pathway (*PAL, C4H*, and *4CL*), which allocate precursors for monolignol synthesis. It is tempting to speculate that supply of precursors limits lignin formation leading to different lignin amounts observed in MsSND1, MsSCM1, and MsMYB103 (Fig. 4A). Similarly, the high expression of several putative *HCT, C3’H*, and *CCoAOMT* genes in MsSND1 and MsSCM1 may channel resources for generation of lignins rich in guaiacyl and syringyl units (Fig. 4B; Fig. 6). Interestingly, *MsMYB103* expression leads to less pronounced increase in the expression of these genes, whereas transcripts for F5H and COMT, which redirect monolignol precursors towards sinapyl alcohol, are induced more strongly than by expression of MsSCM1, possibly accounting for the high accumulation of syringyl units in the cell wall lignin (Fig. 4B). It has been suggested that AtMYB103 specifically regulates F5H expression to modulate lignin S:G ratio (Öhman *et al*., 2013). However, this direct and specific link could not be established for MYB103 proteins originating from grass species, which were shown to regulate cellulose and hemicellulose (Hirano *et al*., 2013; Yang *et al*., 2014; Ye *et al*., 2015). In line with these results, MsMYB103 seems to orchestrate and finetune SCW formation and lignin biosynthesis more broadly. This notion is supported by the distinct expression profiles of various putative laccases and peroxidases (Fig. 6), which may also have implications on lignin composition (Berthet *et al*., 2011; Voxeur *et al*., 2015; Fernández-Pérez *et al*., 2015). Taken together, the transcriptomic data provide a detailed atlas of the transcriptional network of MsSND1, MsSCM1, and MsMYB103 underlying lignin biosynthesis and SCW formation.

**Fig. 6.**
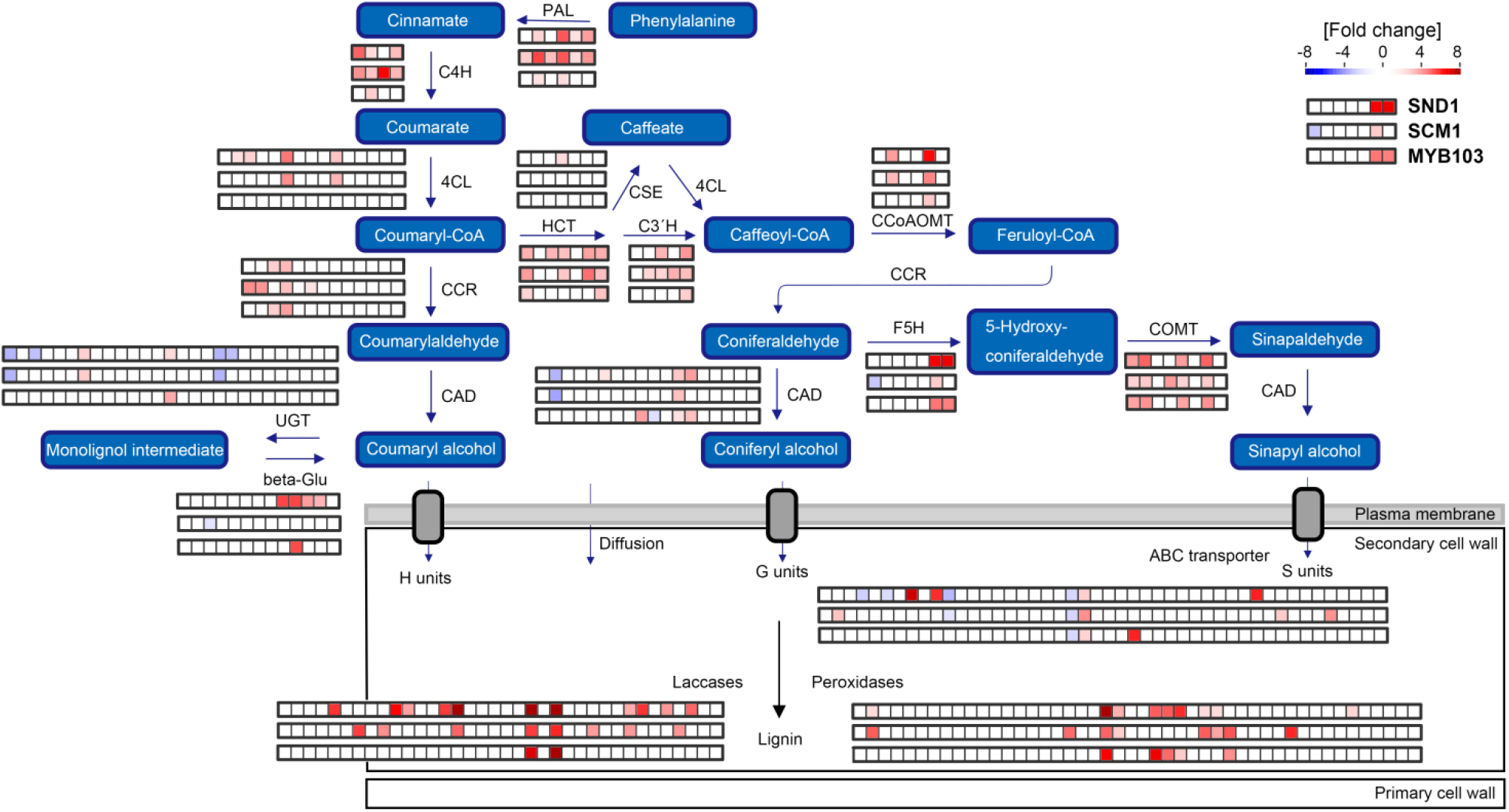
A schematic overview of lignin biosynthesis pathway and DEGs regulated by MsSND1, MsSCM1, and MsMYB103. The fold change of putative genes involved in lignin biosynthesis according to functional annotation by Kourelis *et al*., (2018) are represented in colored squares, generated by MapMan software (Thimm *et al*., 2004). The rows of squares visualize the three samples MsSND1, MsSCM1, and MsMYB103, respectively. Red indicated up-regulation, while blue indicated down-regulation. 4CL, 4-coumarate CoA ligase; beta-Glu, β-Glucosidase; C3’H, *p*-coumaroyl shikimate 3’-hydroxylase; C4H, cinnamate 4-hydroxylase; CAD, cinnamyl alcohol dehydrogenase; CCoAOMT, caffeoyl CoA *O*-methyltransferase; CCR, cinnamoyl CoA reductase; COMT, caffeic acid *O*-methyltransferase; CSE, caffeoyl shikimate esterase; F5H, ferulate 5-hydroxylase; HCT, hydroxycinnamoyl CoA:shikimate hydroxycinnamoyl transferase; PAL, phenylalanine ammonia lyase; and UGT, UDP-Glucuronosyltransferase.

## Conclusion

In order to explore the potential of lignocellulosic biomass as sustainable resources, lignin engineering has become a focus of research with the goal to either decrease recalcitrance or to produce more valuable lignins with enhanced utility. The genus Miscanthus has gained momentum as a prospective lignocellulosic feedstock, but the mechanisms underlying lignin and SCW formation are only poorly investigated. In this study, we identify and characterize three TFs, namely MsSND1, MsSCM1, and MsMYB103 as regulators involved in lignin biosynthesis associated with distinct cell wall compositions that present interesting targets for specific cell wall manipulations. In order to bypass challenging and time-consuming transformation procedures of Miscanthus, infiltration-RNA-seq was investigated as a suitable method to uncover transcriptional regulation of TFs involved in SCW biosynthesis and beyond. The extensive transcriptomic data in response to expression of *MsSND1, MsSCM1*, and *MsMYB103* provided in this study, offer the opportunity to gain a deeper understanding of lignin formation. Despite intense research focusing on the lignin biosynthetic pathway, it is still a matter of debate how monolignols are trafficked to cell wall and which are the driving forces behind this process. It also remains elusive if glycosylated monolignols play an essential role as storage and/or transport molecules. Comparison of the three transcriptomes allowed for the identification of potential candidate genes involved in storage (UGTs, β-Glucosidases), transport (ABC transporter) and/or oxidative polymerization (laccases, peroxidases), providing a valuable starting point for future research.

## Acknowledgements

The authors would like to thank Markus Kiefer (Centre for Organismal Studies, Heidelberg) for providing access to computational resources. This work was supported by a grant from the Ministry of Science, Research and the Arts of Baden-Württemberg (FKZ: 7533-10-5-74) to Thomas Rausch. Research in the laboratory of Sebastian Wolf is supported by the German Research Foundation (DFG) through the Emmy Noether Programme (WO 1660/2). Feng He and Wan Zhang were supported by the Chinese Scholarship Council with fellowship (201506160060 and 201508130065).

## Author contributions

P.G., F.U., S.M., T.R., and S.W. designed the experiments; P.G., E.M., J.X., W.Z. and F.H. performed experiments and analyzed data; P.G., T.R., and S.W. wrote the manuscript.

## Availability of data

The RNA sequencing datasets obtained in this study are available in the Sequence Read Archive (SRA) database under BioProject PRJNA529062 or accessions SRR8786162 - SRR8786173.

---

**Supplementary Fig. S1.**
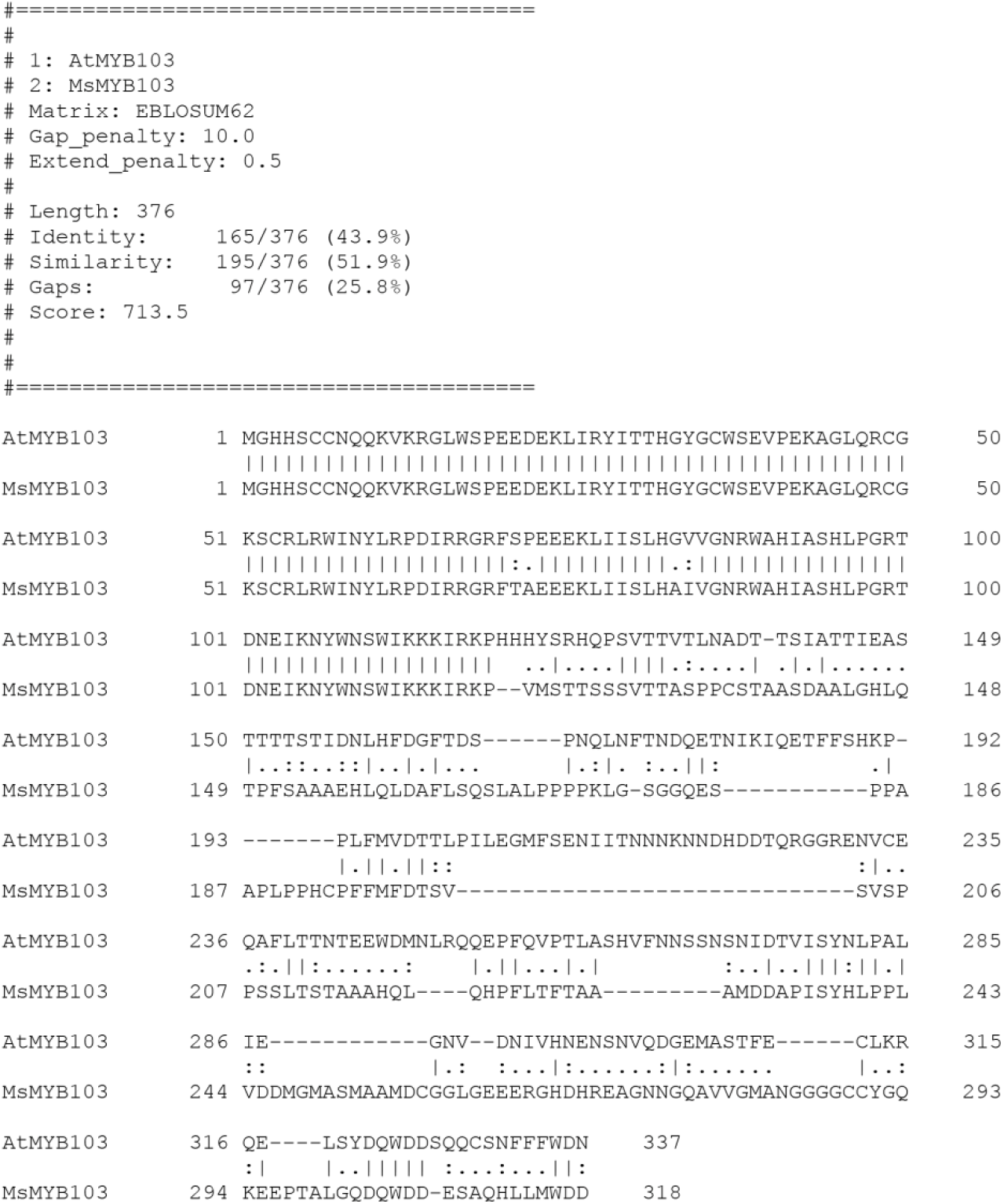
Alignment of AtMYB103 and MsMYB103.

**Supplementary Fig. S2.**
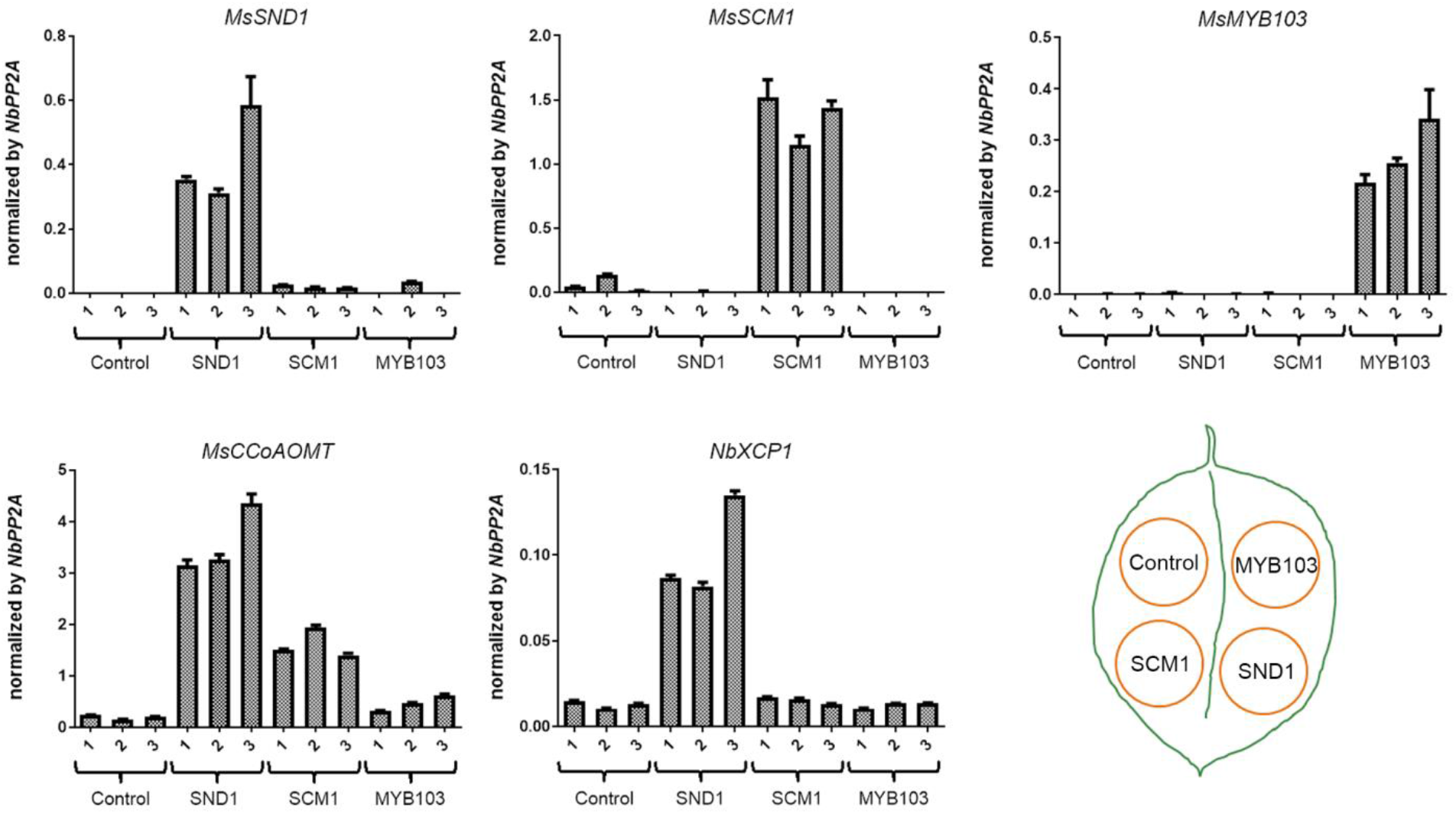
Expression analysis of *MsSND1, MsSCM1*, and *MsMYB103* and two putative targets in tobacco samples used for RNA-seq experiment. Each leaf was infiltrated according to the infiltration scheme to obtain three biological replicates. Four days post infiltration expression of respective gene was determined by q-RT-PCR and normalized against PP2A. Error bars represent + SE.

Supplementary Fig. S3. Detailed heatmap of normalized counts (logCPM) of MsSND1, MsSCM1, and MsMYB103 including biological replicates and gene descriptions and/or gene identifier.

Supplementary Tab. 1. Primer sequences, GreenGate modules, and assembled constructs.

Supplementary Tab. 2. List of DEGs of MsSND1, MsSCM1, MsMYB103, and intersection

Supplementary Tab. 3. GO enrichment analysis of up- and downregulated DEGs following expression of *MsSND1, MsSCM1, MsMYB103*, and intersection.

